# ROSLT: An Improved Raspberry-Pi Open-Source Live Voluntary-Wheel Running Tracker Method and Resource

**DOI:** 10.1101/2024.12.02.626456

**Authors:** Patrick Sanosa, Jade P. Marrow, Amelia R. Malicki, Keith R. Brunt, Jeremy A. Simpson

## Abstract

**Background:** Exercise promotes health and has therapeutic effects on disease. Over time, the body improves its maximal exercise capacity through training adaptations such as an increase in VO2 max. In mice, voluntary wheel running allows for a natural setting to test spontaneous running behaviour under non-stressed conditions. There is a need to design sensitive animal-based assay that improves resolution for differentiating exercise performance from a regular cyclometer (which presents a single value from a summary of dynamic data collected over time) and offer circadian analyses. The purpose of this work is to examine the exercise behaviours of mice with a focus on circadian rhythm of running. We hypothesize the pi cyclometer (ROSLT) will mirror VDO M2.1 behaviors, enhancing circadian rhythm insights.

**Methods:** Using a hand-built cyclometer programmed through the Raspberry Pi computer, voluntary wheel running behaviours in CD-1 mice (∼8-10 weeks) were recorded for 6 consecutive days. This features a Hall Effect sensor and neodymium magnets attached on the running wheels that will detect changes to wheel rotation, speed, acceleration, and distance (continuously) and publish the data to a server in real-time. To compare capabilities, running wheels will also be equipped with the VDO M2.1 WR Cycling Computer to track distance which will be manually recorded once a day. Accuracy from both devices were mechanically validated.

**Results:** The main findings include that voluntary wheel running distance over 6 days produces inaccuracies by the VDO. The VDO showed fluctuations in distance over the last 3 days, while ROSLT showed consistent measurements.

**Conclusion:** This comparison shows that ROSLT expands on the running activity of mice each day while maintaining accuracy and precision. This novel dynamic circadian cyclometer will advance our research abilities and can be used in differentiating exercise performance in applications such as doping.

## Introduction

Exercise is vital to our health span and recovery from injury or disease. Improving maximal exercise capacity by training to increase in VO2 max is one of the central goals for cardiovascular fitness^9^. Exercise research often captures the physiological changes during physical activity, but this requires sensitive and expensive data recording devices.

Current methods for measuring the fitness activity of mice include the Lockable Open-Source Training Wheel^1^, Deitzler and Bira’s Motricity Tracker^2^, Terstege and Epp’s PAW design^3^, Mouse revolution counter MRC^4^, wheel-running activity acquisition WRAQ^5^, DynaLok^6^, and the Open Face Home Cage Running Wheel^7^. All these instrument methods rely on a hall sensor to detect running parameters, such as speed and distance by magnetic field displacement. To expand on the idea of using hall sensors, we envisioned a device that would record speed, distance, and acceleration. Ensuring that a tracker would adapt and record running parameters of any wheel size using live tracking from anywhere in the world and exporting data features. Raspberry Pi is a small programmable computer that stores/transmits data in one system after recording the electrical signal produced by a hall sensor. Moreover, this prevents the need for USBs to collect or transfer data as it is accessible on a cloud server.

Here, we map the development of an inexpensive, efficient, and sensitive device to monitor the real-time fitness and activity of mice in voluntary wheel running conditions aligned with natural circadian behaviours. Accordingly, this is open-source data and equipment, so that any lab can now establish and record voluntary wheel running by using a hall sensor coupled with the Raspberry Pi Model 4B. Data can be stored in a cloud server, allowing for live tracking and user-friendly data exporting. Additionally, the method will be compared to the “gold-standard” VDO M2.1 bike cyclometer that can calculate total distance, average speed, maximum speed, and run time. Comparing our device to the gold standard provides added confidence for researchers regarding accuracy and precision in recording running parameters. Ultimately, this will encourage the development of improved technological assays to measure exercise performance and behaviour in rodents.

## Methods

The Raspberry Pi Model 4B (4 GB RAM with 32 GB micro-SD card) computer was selected to perform the necessary calculations for running distance, speed, and acceleration accurately. The hall sensor was selected as it does not produce any audio/tactile input, which could potentially affect the behaviour of the mice, resulting in an increased running distance. This was previously validated by cite, where they used a regular electromechanical switch to run a mouse, which resulted in a cue for the mice to run at a greater distance^8^. The hall sensors come soldered onto a small circuit board in series with two small green LED lights. One LED is constantly lit while the other only turns on when the hall sensor senses a magnet in proximity. This also permits easier troubleshooting. Based on Seward and Harfmann’s previous experiment, the green LED light does not affect the exercise performance of mice^8^. However, whether the flashing or intensity of the LED light affects mouse running behaviour is still unclear. Neodymium earth magnets were used for the magnets to enhance the sensitivity and detection of a magnetic presence with the hall sensor. To manage multiple hall sensors, a wiring diagram was made on the breadboard along with a small power supply module to handle the appropriate currents running through the sensors to prevent overloading the Raspberry Pi (see Figure 1). Running parameters, such as average acceleration, speed, distance, and rotations were continuously recorded by the Raspberry Pi and uploaded to an online database (IoT service; ThingSpeak) every minute for live data view and exporting (see Figure 3).

**Figure 1.**
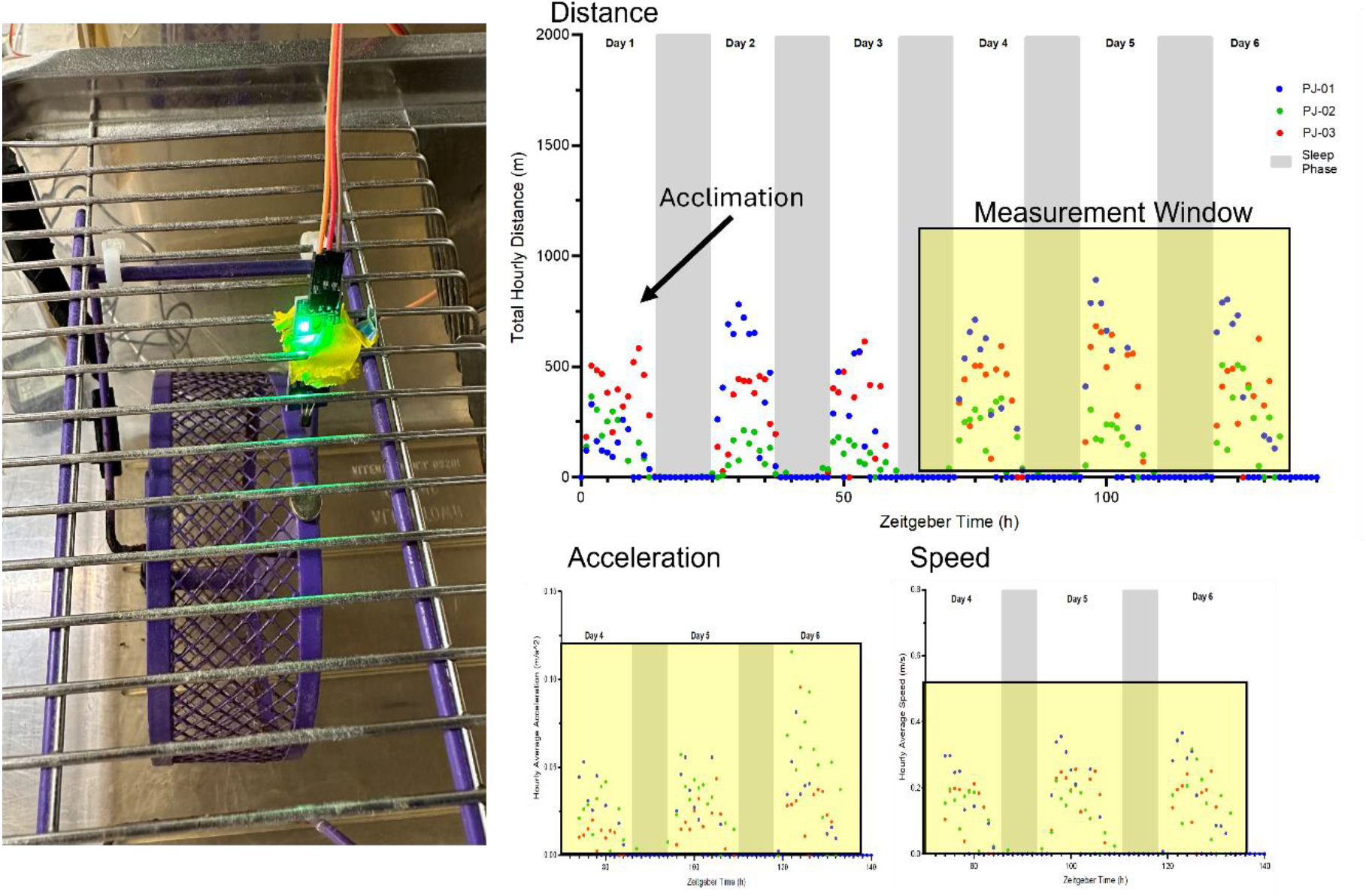
Circadian analysis between distance, acceleration and speed recorded by ROSLT over 6 days. Each point is the average or total of a parameter from each mouse per hour (n = 3). The zeitgeber time corresponds to hours throughout the day starting at 0 = 8 pm, where the mice’s active hours began followed by 1 = 9 pm, 2 = 10 pm, etc. This was plotted using PRISM after the raw data was extracted from ThingSpeak and imported into a pipeline analysis script written in R.

The ROSLT was tested and compared against the VDO M2.1 cyclometer system (the current gold standard; VDO M2.1 cyclometer, Sigma-Elektro GMBH, Neustadt, DE). The setup included the VDO M2.1 sensor placed laterally to the upright running wheel, while the hall sensor was placed above the wheel. The location was chosen for the hall sensor since placing it in the same position as the VDO M2.1 sensor would make it susceptible for the mice to chew (see Figure 1). A major difference between the two methods is the weight of the magnets: the neodymium magnet weighed 1.5 g, whereas the VDO M2.1 magnet weighed 4.1 g. Certainly, this may affect the torque/balance of the wheels and influence its tendency to sway clockwise to counterclockwise when the systems are assessed separately (see Figure 1). To address this issue, video recordings during a mouse’s wake-phase (∼2-3 AM) will demonstrate whether the wheel is concurrently rocking back-and-forth with the mouse (see GitHub repository where assessment with TAPO security camera was performed).

The voluntary wheel running apparatus consisted of a large plastic mouse cage (12cm diameter), with an attached upright running wheel (see Figure 1). The running wheel was equipped with a computerized bicycle monitor (VDO M2.1 WR Cycling Computer) used to measure four parameters: running distance (km), running duration (hh:mm:ss), average speed (km/hr), and maximum speed (km/hr), along with a hall sensor recording the average speeds, acceleration, distances, and rotations every minute across the 24 hours of a day. However, on the cycling computer, the parameters were recorded once every 24 hours, and the monitors were reset daily. This cyclometer was used to compare and assess the accuracy of ROSLT. The hall sensors were placed on top of their respective cage with the appropriate wiring to the Raspberry Pi. The VDO M2.1 cycling computer’s magnet was attached to the side of the wheel opposite the magnet that will be detected by the hall sensor. This specific setup is designed to minimize the amount of torque/imbalance of the wheel (see Figure 2). The VDO M2.1 monitors were reset daily, and recordings were logged for 6 consecutive days (5 nights), while ROSLT continuously tracked and published the data to a live server (ThingSpeak) on a minute-to-minute basis (see Figure 3).

**Figure 2.**
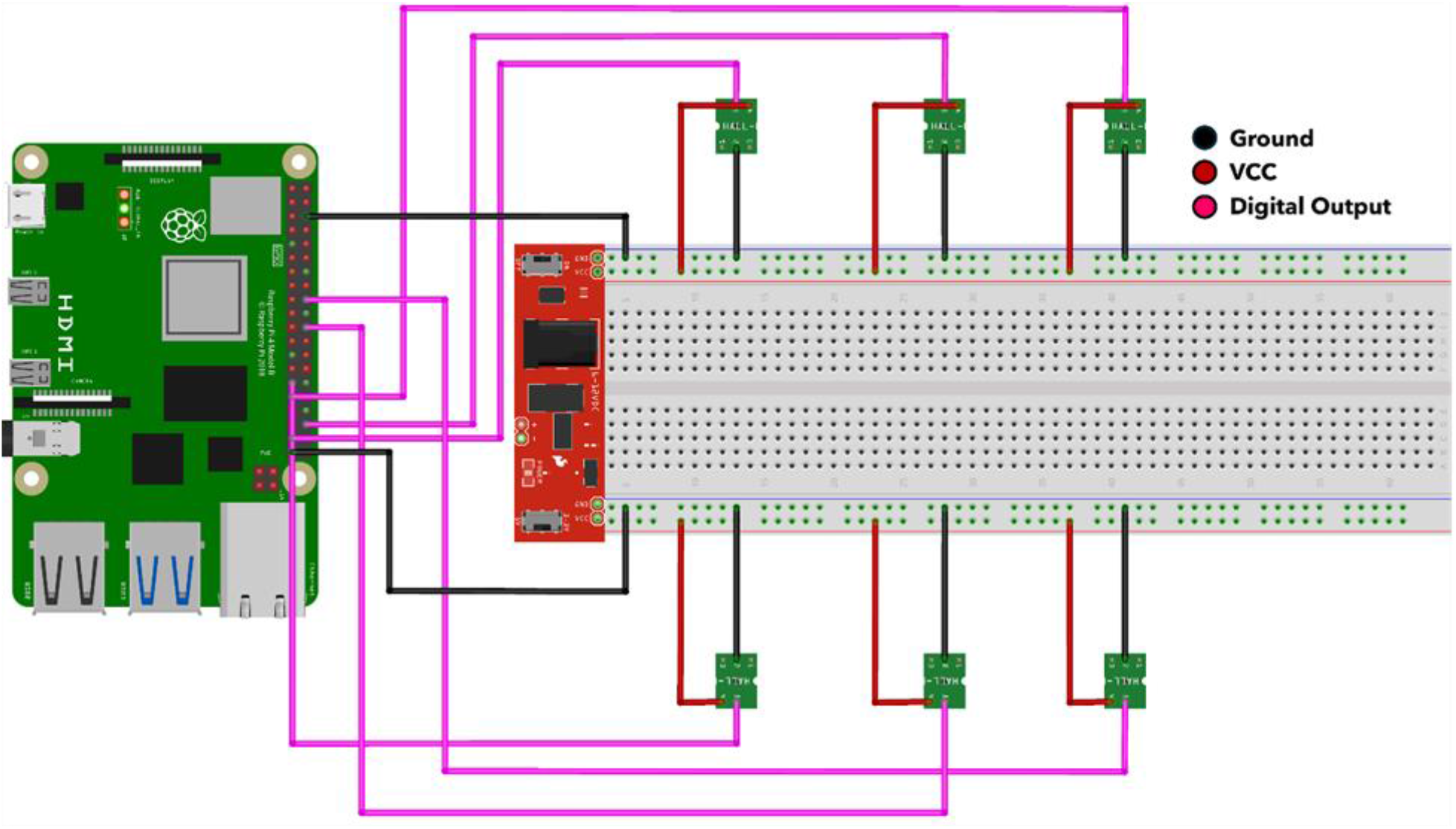
Schematic diagram of wiring connection between the hall sensors and the Raspberry Pi. The black and red wires represent the ground and supply voltage, respectively. The pink-coloured wires relay the signal (digital output) to the Raspberry Pi from each sensor. All sensors were powered by the 5V supply from the breadboard power supply that used a 12V power adapter. Additional sensors were added to the diagram to serve as an example for the expanding capabilities of ROSLT.

**Figure 3.**
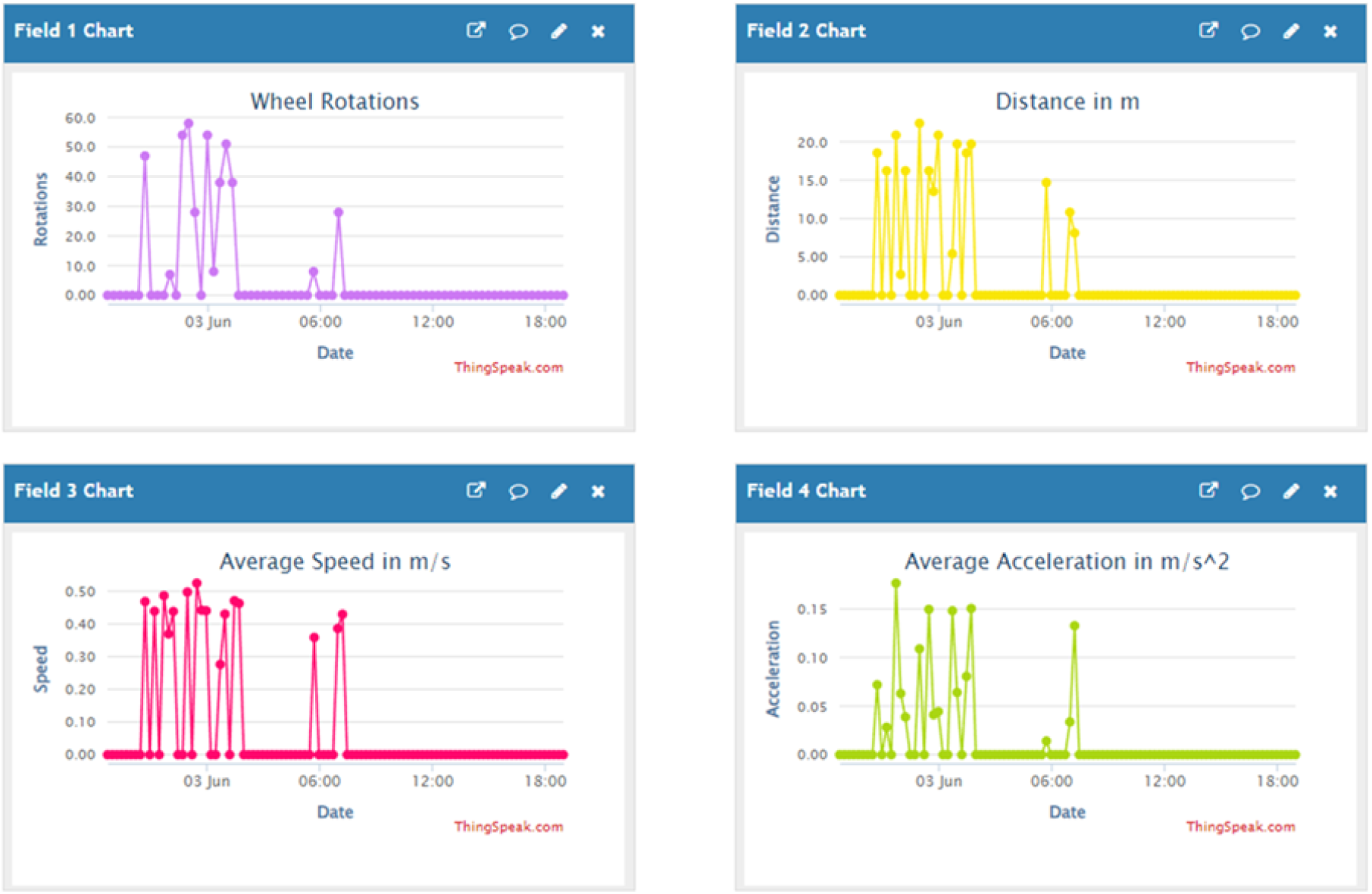
This showcases a live-data server to track real-time running parameters for a rodent. This uses the IoT service from ThingSpeak. The data can be exported as a CSV file.

### Validation of ROSLT and VDO M2.1 Cyclometer simultaneously in vivo

All mice used were 7–8-week-old CD1 males n = 3 (Charles River Laboratory International Inc.). As it was not hypothesis testing, we elected to use one sex only. Validation of the VDO M2.1 cyclometer was assessed through 3 parameters: the total distance, speed, and run time. All 3 cages were assessed across 5 nights. Mice were put in their cages at 5 PM (3 hours before the dark cycle of the mice) and recording ended at 5 PM in the afternoon on the 5th day. Mice were housed at 23-24C with 45% humidity and maintained on a 12h light/dark cycle with food and water provided ad libitum. This study was approved by the Animal Care Committee at the University of Guelph and all experiments were conducted in accordance with the guidelines from the Canadian Council on Animal Care.

### Mechanical validation of ROSLT and VDO M2.1

A 12 V DC motor (from an electric toothbrush) was used with the speed of the motor controlled by a PWM board (pulse width modulation), which was supplied through the Raspberry Pi’s 5V pins. The validation of ROSLT and the VDO M2.1 was assessed by two means: The ROSLT’s sensor placed by itself adjacent to a foam wheel and recorded for three separate sessions (for a total of 6 hours) in various speeds with a wheel circumference input of 79 mm, and both the VDO M2.1 and ROSLT’s sensors placed adjacent to a foam wheel with a wheel circumference input of 386 mm and stopped at a set distance.

### Data Analysis

After the raw data from ThingSpeak was exported as a CSV (comma-separated values) file, it was processed through an R-script that automatically organizes the data to get SEM, mean, and total values per hour for import into PRISM. The open-source code and raw data can be found on our GitHub repository [insert link here]. Analysis for in vivo experiments was done using R version 4.3.2 (R Core Team, 2020) packages. Graphing and statistical analyses were performed using Prism 10 statistical software (GraphPad Software, San Diego, CA).

## Results

### Mechanical and in-vivo validation of ROSLT and VDO M2.1 Cyclometer

All 3 male CD1 mice (∼ 7-8 weeks old) were placed in their cages and ran for 5 nights. To further test for accuracy based on location differences of the hall sensor, we decided to validate this using mechanical methods (shown below).

## Discussion

### ROSLT is accurate through mechanical validation

The first assessment used a 12 V DC motor that ran for a total of 6 hours divided into three separate sessions using ROSLT (Table 1). The total distance calculated from the fixed speeds and duration were as follows: Run 1 at 1080 m, run 2 at 4320 m, and run 3 at 9720 m. In contrast, ROSLT recorded values at approximately 1036 m, 4225 m, and 9543 m, respectively. As expected, ROSLT recorded the average speeds at 0. 297, 0.6, and 0.9 m/s, respectively. The percent error difference between the calculated values and ROSLT’s recorded values for all three parameters were less than 5%. This demonstrates that ROSLT can accurately record distance and speed while uploading live data to produce minute-by-minute changes in running behavior.

**Table 1.** Accuracy assessment of ROSLT from its recorded run distance and speed to hand-calculated values. The three sessions were fixed across three different speeds for 1-3 hours and ran with a DC motor which included a speed control. The hand-calculated values come from the fixed values of speed and duration the motor ran. The hall sensor was placed directly lateral to the disc on the motor. The percentage error difference was within ∼0-4%.

| Run Variables (79 mm disc circumference) | ROSLT Values (Total, Mean, SEM) | Calculated Values | % Error Difference |
| --- | --- | --- | --- |
| **Distance (m)** |  |  |  |
| Run 1 (1 hour, x1 speed) | 1036.6; 17.9 +/- 0.020 | 1080 | 4.1 |
| Run 2 (2 hours, x2 speed) | 4225.2; 36.1 +/- 0.025 | 4320 | 2.2 |
| Run 3 (3 hours, x3 speed) | 9543.4; 53.9 +/- 0.111 | 9720 | 1.8 |
| Mean +/- SEM |  |  | 2.7 +/- 0.71 |
| **Average Speed (m/s)** |  |  |  |
| Run 1 (1 hour, x1 speed) | 0.297 +/- 0.0003 | 0.3 | 1.0 |
| Run 2 (2 hours, x2 speed) | 0.601 +/- 0.0004 | 0.6 | 0.14 |
| Run 3 (3 hours, x3 speed) | 0.901 +/- 0.0009 | 0.9 | 0.14 |
| Mean +/- SEM |  |  | 0.43 +/- 0.29 |

### ROSLT is consistent and accurate in-vivo

Using ROSLT simultaneously with the VDO M2.1, three mice were recorded across 5 nights then exported from ThingSpeak into RStudio (using an R-script) to organize the data parameters into averages and totals each hour. This was then imported into PRISM to get a visual on the circadian rhythm of the mice (Figure 1). As expected, the mice were most active during the dark hours, where total running distances increased. Additionally, the total running distances gradually decreased during daylight hours. The average speeds and accelerations were expected to be directly proportional to distance, where a downward trend exists when mice ran less (Figure 1). There were also outliers on the ROSLTs recorded accelerations, we suspect that this may be due to the magnet on the wheel rapidly rocking back-and-forth on the hall sensor due to torque/imbalance. Moreover, ROSLT’S shows its reliability to track the fitness activity and the circadian behaviour of mice.

To test the accuracy of ROSLT, we decided to compare the daily total distances ran by each mouse recorded by the VDO M2.1 cyclometer (Figure 4). We decided to emphasize on the last 3 days, since the mice required time to acclimate to their running cages (Figure 1). The fluctuations of the distances recorded by the VDO indicate that values become more inconsistent as mice ran for a longer period of time. However, for ROSLT, the values recorded are similar to the following day, demonstrating that the device is accurate. One explanation for the inconsistent recordings of the VDO device could be that it is not able to calculate wheel rotations accurately when the wheel circumference input is less than an average bike wheel (∼2m).

**Figure 4.**
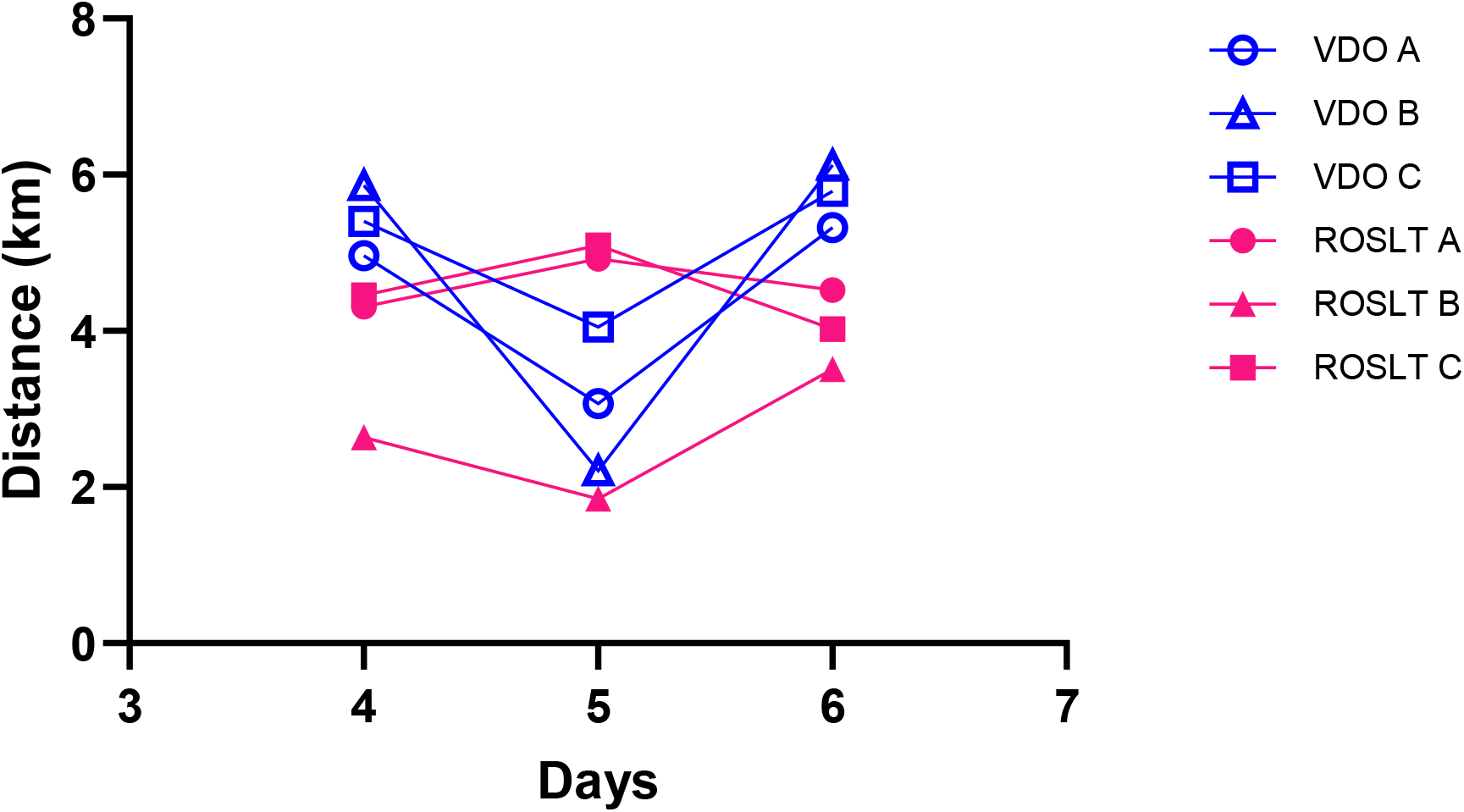
A comparison of the total run distances recorded between the VDO M2.1 Cyclometer and ROSLT on the last 3 days of voluntary wheel running over 6 days. Each point is the total distances ran by each male CD1 mice (7-8 weeks old; n = 3).

### ROSLT is both accurate in-vivo and through mechanical validation

Lastly, to validate ROSLT’s accuracy, a mouse was ran for 24 hours along with the VDO M2.1. The total distance of the mouse recorded by the VDO was half of the ROSLT’s distance recorded (Table 2a). Since the difference between distances was significantly greater with the VDO M2.1, we expected the speed to increase. To further validate ROSLT’s accuracy over 24 hours, we set an absolute distance of 3770 m for a motor to run (see table 2b). The VDO’s inaccurate recording shows that it is unable to deal with circumferences that are smaller than an average bike wheel. Taken together, both the mechanical and in-vivo test for ROSLT show that it is a more precise and accurate method to use when tracking the fitness behaviour in mice.

**Table 2a (mouse).** Accuracy assessment of ROSLT compared to the VDO M2.1 in-vivo over 24 hours (n = 1). Both the VDO and ROSLT sensors were placed vertically above the wheel. The wheel circumference input was 386 mm.

| Parameters | ROSLT | VDO M2.1 Cyclometer |
| --- | --- | --- |
| Total Distance Ran for 24 hours | 3770 m | 7560 m |
| Average Speed for 24 hours | 0.08 m/s | 0.56 m/s |

## Conclusions

In this paper, we present a globally accessible device that can live-track the fitness activity of mice using a Raspberry Pi computer. This device is non-invasive, low-cost, and allows a user-friendly setup for research laboratories. To record a rodent’s circadian behaviour during voluntary wheel running, researchers would be required to purchase a consumer rodent tracker, which could cost up to ∼$1000 for a single cage. (Table 3). Alternatively, the ROSLT device provides research laboratories with a cost-effective and customizable method of tracking the circadian rhythms of mice. This tracker differs from the others due to the addition of two features: an open-source code with live tracking and recording accelerations (Table 3). To validate the accuracy of this tracker, we compared it to a gold-standard, VDO M2.1 bike cyclometer that was previously used in many laboratory settings. We performed various experiments to prove the inconsistencies between the VDO and ROSLT’s recorded parameters. Our findings suggest that the ROSLT can replace the VDO M2.1 Cyclometer as the gold standard, as it is a cheaper and a more sensitive alternative to tracking a rodent’s live exercise activity and circadian rhythm.

**Table 2b (motor).** Accuracy assessment of ROSLT and its absolute distance values compared to the VDO M2.1 cyclometer. A motor was used and set for 3770 m. Both the VDO and ROSLT sensors were placed laterally to the wheel. The wheel circumference input was 386 mm.

| Parameters | ROSLT | VDO M2.1 Cyclometer |
| --- | --- | --- |
| Absolute Distance | 3770 m | 4000 m |
| Measured Speed | 2.5 m/s | 2.8 m/s |

**Table 3.**
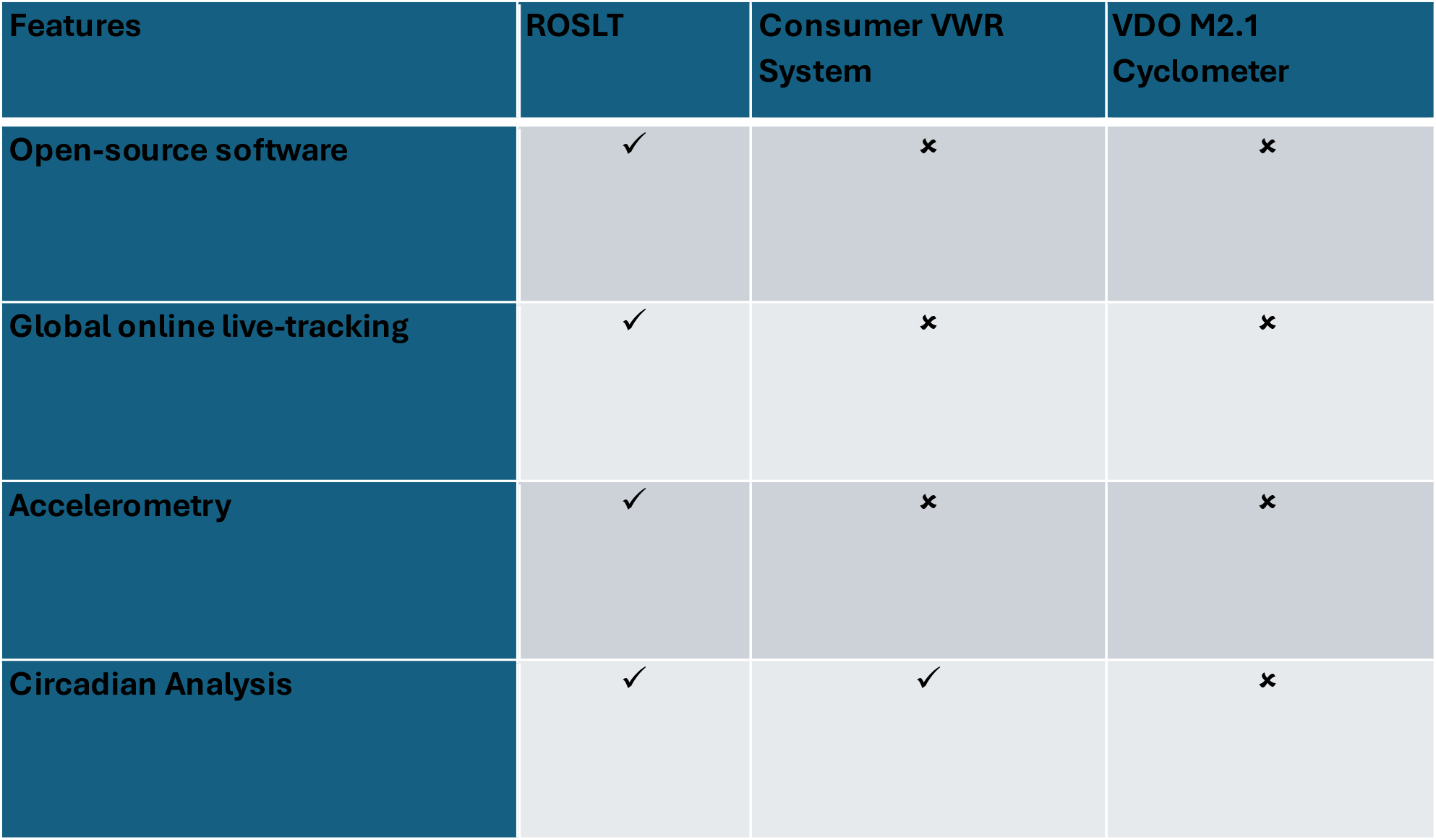
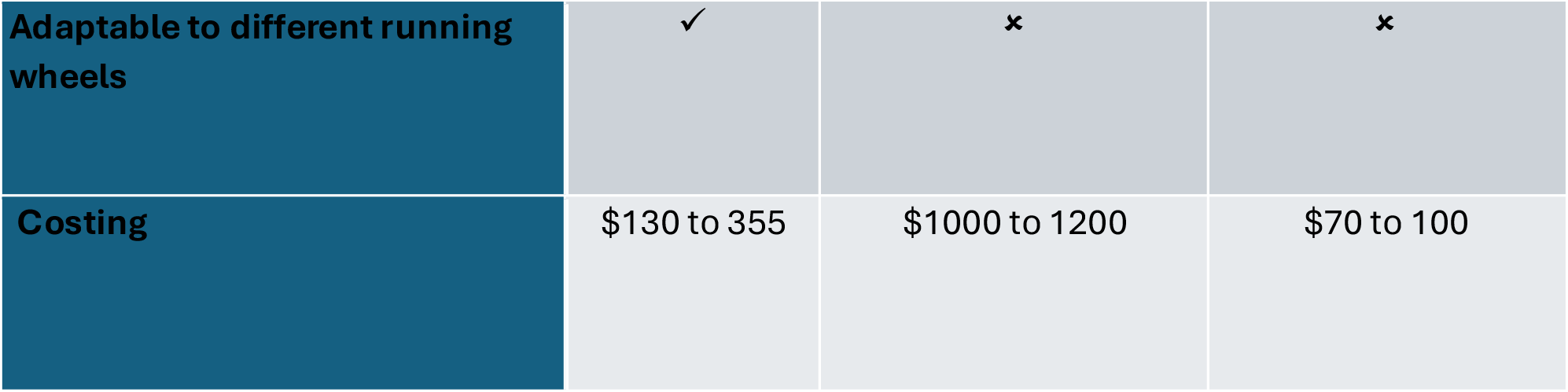
A table comparison between different voluntary wheel running recording methods and their features. Costing is calculated by material and labour/shipping costs.

| Features | ROSLT | Consumer VWR System | VDO M2.1 Cyclometer |
| --- | --- | --- | --- |
| Open-source software | ✓ | ✗ | ✗ |
| Global online live-tracking | ✓ | ✗ | ✗ |
| Accelerometry | ✓ | ✗ | ✗ |
| Circadian Analysis | ✓ | ✓ | ✗ |
| **Adaptable to different running wheels** | ✓ | ✗ | ✗ |
| **Costing** | \$130 to 355 | \$1000 to 1200 | \$70 to 100 |

